# Insertion of fluorescent proteins near the plug domain of MotB generates functional stator complex

**DOI:** 10.1101/2024.09.27.615325

**Authors:** Jyoti P Gurung, Pietro Ridone, Anaïs Biquet-Bisquert, Gary Bryant, Francesco Pedaci, Ashley L. Nord, Matthew AB Baker

## Abstract

Many bacteria swim by the rotation of the bacterial flagellar motor (BFM). The BFM is powered by proton translocation across the inner membrane through the hetero-heptameric MotA_5_MotB_2_ protein complex. Two periplasmic domains of MotB are critical in activating BFM rotation: (1) the peptidoglycan binding (PGB) domain that anchors MotB in the peptidoglycan layer and (2) the plug domain that modulates the proton flow. Existing cytoplasmic fluorescent probes have been shown to negatively affect motor rotation and switching. Here we inserted a fluorescent probe in the periplasm near the plug of MotB in an attempt to circumvent issues with cytoplasmic probes and for possible use in observing the mechanism of plug-based regulation of proton flow. We inserted green fluorescent protein (GFP) and iLOV, a fluorescent version of the light-oxygen-voltage (LOV) domain, in four periplasmic locations in MotB. Insertions near the plug retained motility but showed limited fluorescence for both fluorophores. Additional short, flexible glycine-serine (GS) linkers improved motility but did not improve brightness. Further optimization is necessary to improve the fluorescence of these periplasmic probes.

## Introduction

The bacterial flagellar motor (BFM) is an intricate molecular machine responsible for motility in many bacteria, driving the rotation of flagella to propel cells through their environment. The BFM is driven by various ions such as H^+^ and Na^+^, with the MotA_5_MotB_2_ stator complex specifically utilizing proton flow to generate torque through its interactions with the rotor, facilitating the rotation of the motor (Deme et al., 2020; Santiveri et al., 2020). The N-terminal of the MotB monomer has a small cytoplasmic tail, then a single transmembrane helix in the inner membrane followed by short helix (the plug domain), and an unstructured linker which is finally connected to a peptidoglycan binding (PGB) C-terminal domain (**Figure 1**). It is hypothesised that upon MotB binding to the peptidoglycan layer, the helical structure of the ‘plug domain’ releases from the periplasmic surface of MotA, exposing the ion pore and facilitating the route for H^+^ ion translocation (Hu et al., 2023). The PGB domain is critical in assembling the stator complex around the BFM rotor by anchoring it to the peptidoglycan layer. The stator is dynamic and the off rate of stator binding is modulated in response to increased load (Lele et al., 2013; Nord et al., 2017).

**Figure 1.**
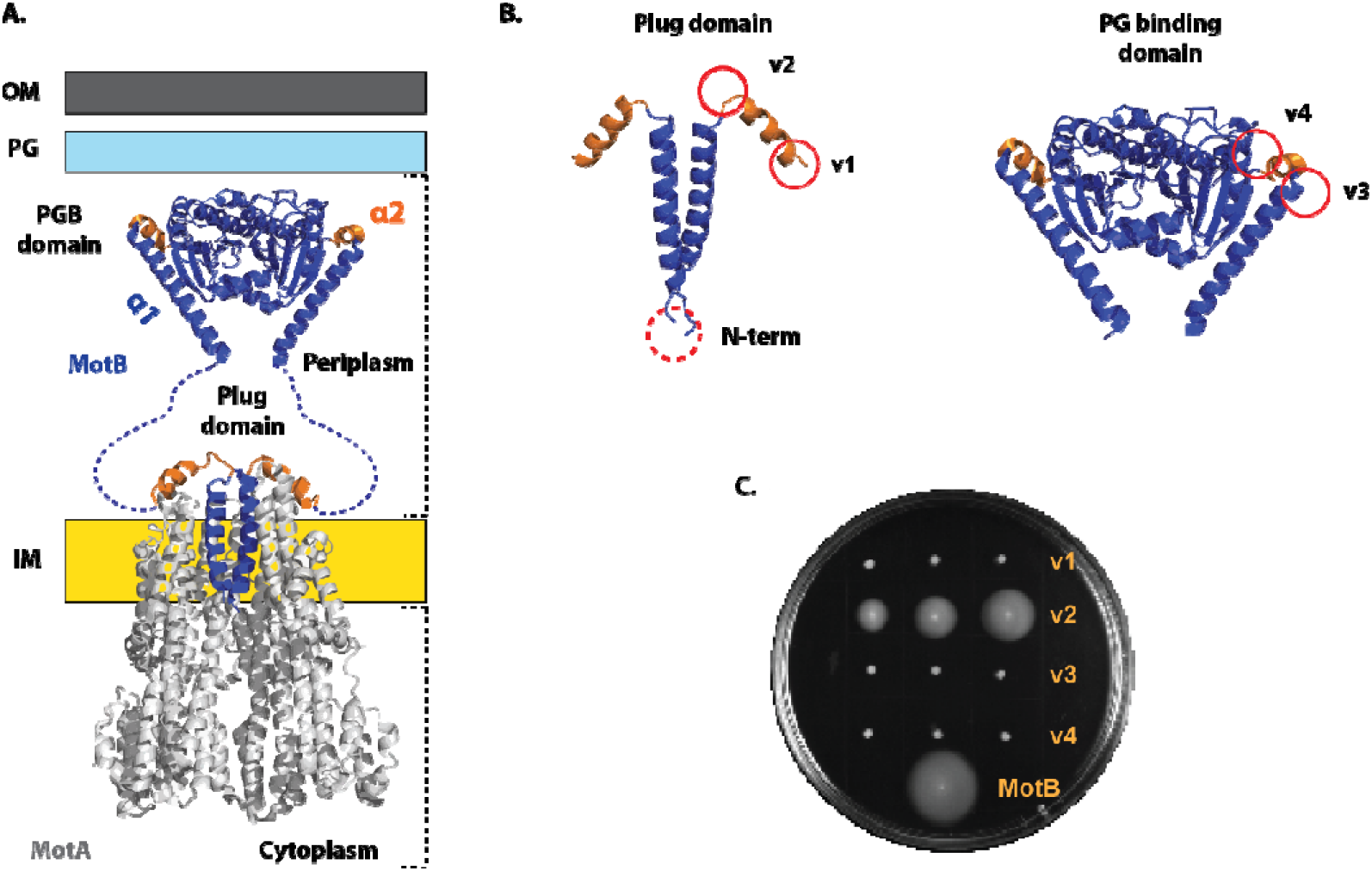
LOV insertion sites in MotB and its impact on bacterial motility. (A) Structure of MotA_5_MotB_2_ stator complex in the bacterial cell membrane. Cryo-EM structure of the stator complex containing MotA (grey-colored) and MotB (blue-colored) with PG binding domain from Salmonella enterica (PDBID: 2zvz): and plug and transmembrane (TM) domain from Campylobacter jejuni (PDBID: 6ykm). (B) Location of LOV domain in MotB (denoted by the red solid circle). v1 and v2 represented the insertion of the LOV domain before and after the plug domain (orange-colored) respectively. Insertion location in the N-terminal of MotB is denoted by ‘N-term’ (dotted red circle). v3 and v4 represented the insertion of the LOV domain before and after the short α2-helix of the PG Binding domain (orange-colored) respectively. (C) Image of swim plates, containing semi-solid media after 24 hours of incubation at 30°C. Bacterial strains: MotB tagged with LOV (v1, v2, v3, v4), and wild-type motile strain as a positive control (MotA_5_MotB_2_ expressing bacterial strain). Three colonies for each bacterial strain were inoculated in a swim plate (except for wild type).

Fluorescent protein labelling can offer insight into single molecule binding events of the stator complex. To date, standard fluorescent proteins such as GFP, which are relatively large in size (∼27 kDa), have been inserted in the N-terminal of MotB (i.e., cytoplasmic labels) (Leake et al., 2006; Tipping et al., 2013) and have been shown to impact the rotational speed and switching of the flagellar motor (Heo et al., 2017). In our work, we selected a fluorescent version of the light-oxygen-voltage (LOV) domain i.e., iLOV (improved LOV) protein as a small periplasmic fluorescent probe for MotB (Christie et al., 2012). The iLOV fluorescent protein has an advantage over standard fluorescent proteins such as GFP due to its size (∼11 kDa), its oxygen-independence (it can fluoresce in anaerobic conditions) and functionality under a broad range of pH (4-11) (Drepper et al., 2007; Swartz et al., 2001).

The precursor of iLOV, i.e. LOV, is a photo-responsive domain containing flavin mononucleotide as a co-factor or chromophore for photo-sensitive activity. LOV domains are present in many kingdoms of life, predominantly in bacteria and plants with diverse functionalities (Crosson et al., 2003). The LOV domain is composed of three motifs: an N-terminal short helix (A’-α-helix), a middle LOV core (β-sheets and α-helix) and a C-terminal long helix (J-α-helix). FMN is bound to the LOV core in a dark state during which J-α-helix is sequestered into the LOV core. Upon blue light exposure, helical structure of J-α-helix unwinds and causes its release from the LOV core to a disordered or distended form. This conformational change of the J-α-helix upon light exposure can be used as an optogenetic tool, for example, the J-α-helix of the LOV domain was used to regulate the opening and closing of potassium channels for neuron function (Jerng et al., 2021).

Here we inserted a photoactivable LOV domain and a fluorescent LOV domain (iLOV) at two sites near the plug domain and two sites near the PGB domain. First, we tested the effect of blue light illumination on bacterial motility when the photoactivable LOV domain was inserted into MotB. We hypothesised that the conformational change in the LOV domain, induced by blue light, would affect motility in a measurable manner. We then tested the motility and fluorescence of fluorescent iLOV that was inserted in MotB to develop it as a potential periplasmic fluorescent probe. We hypothesised that the motility of cells powered by an iLOV-tagged MotB stator would be comparable to wild-type, with minimal disruption in motility compared with the GFP counterpart. Finally, in an attempt to improve the weak fluorescence of iLOV compared to GFP (Mukherjee et al., 2013), we inserted short and flexible linkers in between fluorescent iLOV and MotB periplasmic domains to determine if linkers could rescue fluorescence while retaining bacterial motility.

## Results

### The LOV domain inserted after the plug domain of MotB does not disrupt the bacterial motility

Four sites were selected in the periplasmic region of MotB for the insertion of photoactivable LOV domain. Two were near the plug domain; one before the α-helical plug at the 50^th^ amino acid of MotB (henceforth known as ‘v1’), and one after the plug domain at the 64^th^ amino acid of MotB (henceforth known as ‘v2’). The two remaining sites were before the PGB at the 128^th^ amino acid of MotB (henceforth ‘v3’) and after the PGB at the 187^th^ amino acid of MotB (henceforth ‘v4’) (**Supplementary Fig. 1**). As a result, four periplasmic variants (v1, v2, v3, and v4) were constructed and the effect on bacterial motility was tested (**Figure 1.B)**. There was no impact on cell growth for all the bacterial strains we prepared (**Supplementary Fig. 2**). When these strains were tested ion semi-solid media (swim plates), periplasmic variant (v2 i.e., LOV inserted after the plug domain) spread measurably after 24 hours of incubation, whereas v1, v3, and v4 did not (**Figure 1.C**). Furthermore, motility in swim plates exhibited by periplasmic variant (v2) and other non-motile variants was verified in liquid media by measuring the swimming velocity using differential dynamic microscopy (DDM) (**Supplementary Table 1 & Supplementary Fig. 3**). When illuminated by blue light (465 nm, 150 lx), no notable differences in swim diameter on swim plates were observed for all variants compared with dark conditions (**Supplementary Fig. 3)**. After 48 hours, v3 and v4 regained motility (**Supplementary Fig. 4 & 5)**, however this was confirmed to be due to restored fully functional MotB from the reversion of a stop codon in the ΔmotB strain (RP3087) (**Supplementary Fig. 6**).

### MotB tagged with iLOV in the cytoplasmic (N-term) and periplasmic variants (v2) spread further on a swim plate than the GFP counterpart

As blue light showed no effect on motility when the photo-switchable LOV domain was inserted, we chose the periplasmic variant (v2 i.e., insertion site after the plug domain) for further development as a periplasmic fluorescent sensor since it had swimming behaviour close to wild-type. We replaced the LOV domain in the v2 variant with PhiLOV2.9 protein, an improved photostable version of the fluorescent iLOV domain (Christie et al., 2012). To assess the impact of iLOV and GFP insertions on bacterial motility, and their fluorescence intensity and utility, we inserted these fluorescent proteins at cytoplasmic end (N-terminal) and periplasmic site (v2) (**Figure 2.A**).

**Figure 2.**
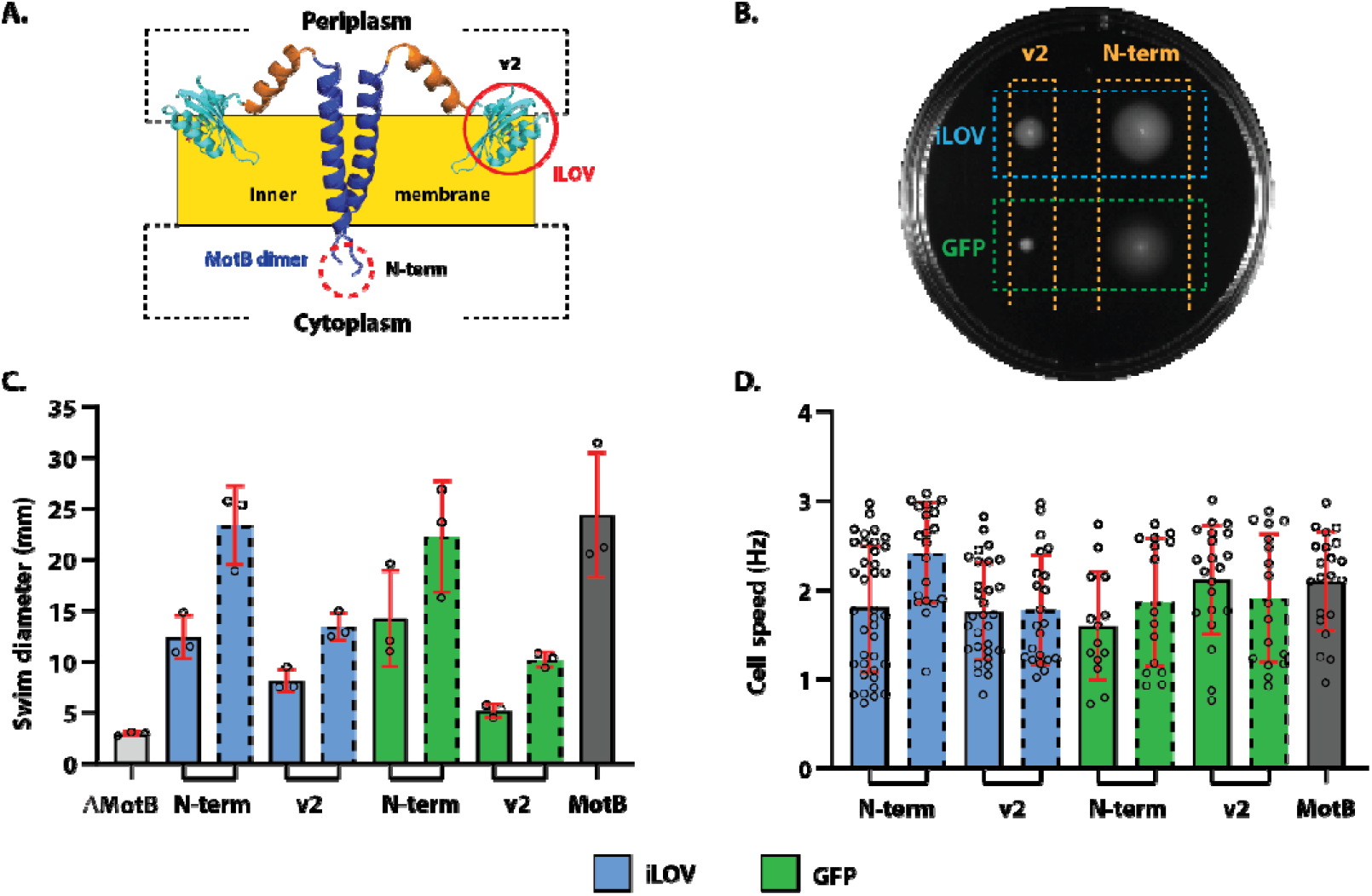
Motility assay for bacterial strains with fluorescently tagged MotB. (A) Schematic diagram of iLOV inserted in the periplasmic domain of MotB at the v2 site. (B) Image of swim plate showing bacterial strains after incubating for 24 hours at 30°C. (C) Bar graph of swim ring diameter (mean ± standard deviation, triplicates data) measured for the bacterial strains tested. Dotted lines indicate strains with linkers. (D) Bar graph of rotational speed (mean ± standard deviation) of tethered cells for each strain. Number of cells (n) averaged to determine mean rotational speed are listed in **Supplementary Table 2**. The bacterial strains tested include ΔmotB (motB deleted strain), GFP-MotB (MotB with GFP tag at the N-terminus), iLOV-MotB (MotB with ILOV tag at the N-terminus), MotB-GFP-v2 (MotB with GFP tag after the plug domain), and MotB-iLOV-v2 (MotB with ILOV tag after the plug domain).

In periplasmic variant (v2), iLOV-tagged MotB showed swimming diameter of 8.1 ± 1.1 mm that was greater than its GFP counterpart (5.2 ± 0.7 mm), but, smaller than wild type (24.4 ± 6.1 mm) (**Figure 2.B** and **Figure 2.C**). However, the cytoplasmic variant (N-term) showed greater motility than the periplasmic variant (v2). The introduction of GS linkers further improved bacterial motility in all variants (**Figure 2.C**). For the cytoplasmic variant (N-term), swim diameter of GFP-tagged MotB increased from 14.2 ± 4.7 mm to 22.3 ± 5.5 mm, whereas swim diameter of iLOV-tagged MotB increased from 12.4 ± 2.1 mm to 23.4 ± 3.9 mm (**Figure 2.C**). Similarly, for the periplasmic variant (v2), a linker increased the swim diameter of GFP-tagged MotB from 5.2 ± 0.7 mm to 10.2 ± 0.7 mm, whereas for the iLOV-tagged MotB a linker increased the swim diameter from 8.1 ± 1.1 mm to 13.4 ± 1.3 mm (**Figure 2.C**). We did not observe any statistically significant difference (with p-value > 0.05) in the average rotational speed between bacterial strains with and without GS linker using a tethered cell assay (**Figure 2.D**).

### Comparative rotation of MotB tagged with iLOV with MotB tagged with GFP for periplasmic (v2) and cytoplasmic variants (N-term)

To assess the rotational speed and switching of the motor powered by our MotB variants we attached 1.1 µm polystyrene beads onto truncated and hydrophobic (FliC_sticky_) bacterial filaments (***Figure 3***) and tracked their rotation. In the cytoplasmic variant (N-term), we observed that the direct fusion of GFP in the N-terminal of MotB reduced the motor speed to 37.4 ± 11.8 Hz compared to the wild type (48.2 ± 15.8 Hz) which was in agreement with previous measurements (Heo et al., 2017). Direct fusion of iLOV in the N-terminal of MotB showed motor speed of 44.4 ± 9.1 Hz, more than the GFP counterpart but slightly less than the wild type. Inserting a short and flexible GS linker in between iLOV and the N-terminal of MotB decreased the motor speed to 26.4 ± 13.8 Hz. In the periplasmic variant (v2), the motor speed of iLOV-tagged MotB was 38.2 ± 13.8 Hz, higher than the GFP counterpart (19.4 ± 7.8 Hz). As per the cytoplasmic variant, insertion of GS linkers did not increase the motor speed for the periplasmic variants (v2) but decreased the speed to 24.6 ± 13.0 Hz. For all the constructs, the speed both counterclockwise (CCW) and clockwise (CW) was lower than WT rotational speed (***Figure 3***).

**Figure 3.**
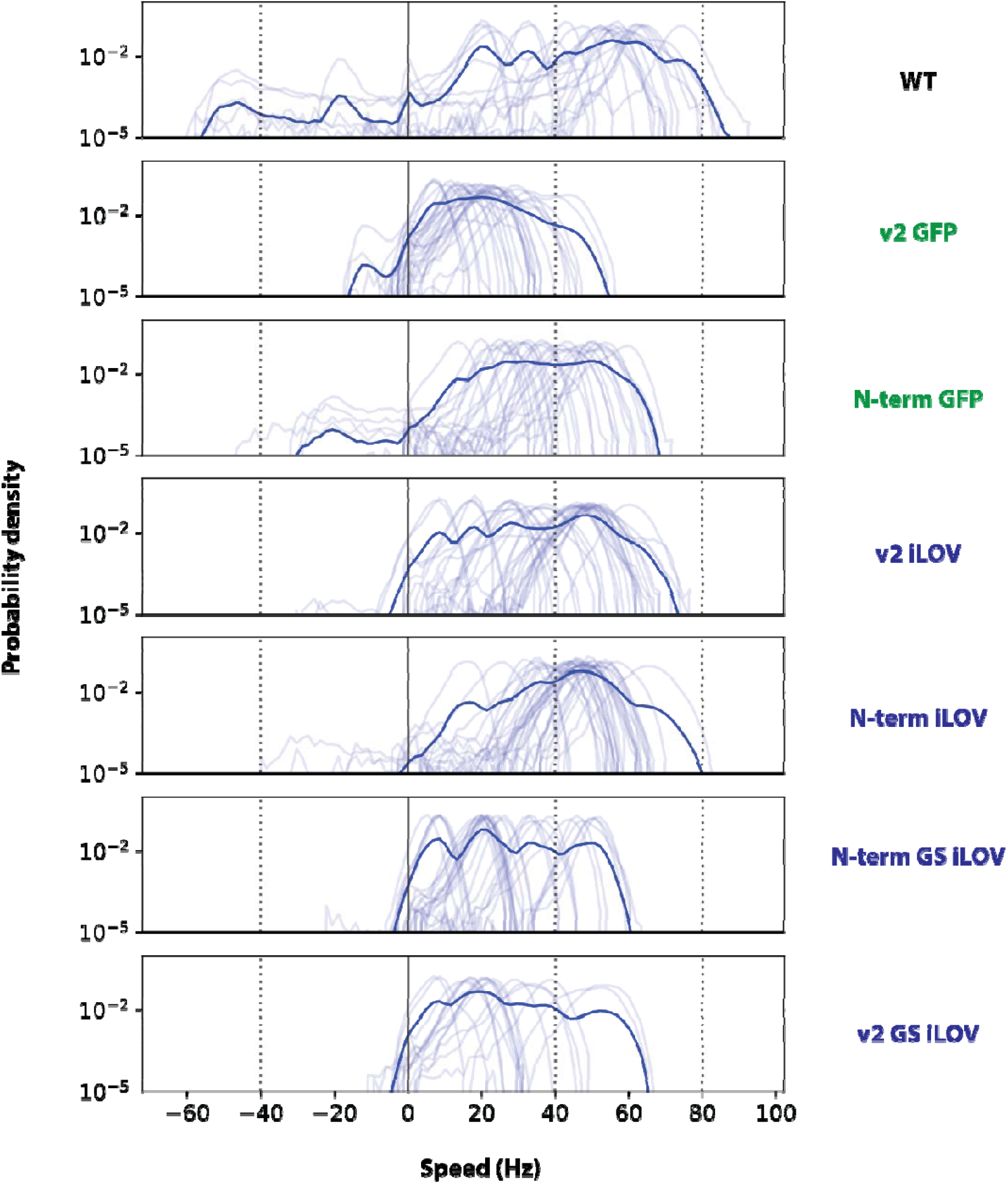
Probability density of CCW-biased motor speeds of iLOV and GFP tagged MotB in N-terminal and v2 position, measured using bead assay. Positive and negative values of speed represent motor rotating in CCW and CW direction, respectively. Light blue lines indicate single motor speed probability densities and thick solid blue line the mean probability density. Number of motors measured: (**Supplementary Fig. 7**): ‘n’ = 23 for WT, 29 for v2 GFP, 27 for N-terminal GFP, 34 for v2-iLOV, 34 for N-terminal iLOV, 30 for N-terminal iLOV GS, and 17 for v2 iLOV GS. Here, ‘GS’ indicates the linker with two amino acids of Glycine-Serine.

Compared to wild type, both cytoplasmic (N-term) and periplasmic variants (v2) showed reduced switching frequency, as seen previously (Heo et al., 2017) (**Supplementary Fig. 8**). The switching frequency of iLOV-tagged MotB in the periplasmic variant (v2) was reduced compared with the GFP counterpart. Insertion of GS linkers in iLOV variants increased the switching frequency, most notably in the periplasmic variant (v2), albeit not to WT levels (**Supplementary Fig. 8**).

### Both GFP and iLOV show limited fluorescence in the periplasm compared with the cytoplasm

We measured the fluorescence for both GFP and iLOV-tagged MotB by comparing the maximum absolute pixel intensity inside a region of interest around a selected cell **(Figure 4)**. In the cytoplasmic variant (N-term) both iLOV-tagged MotB and GFP-tagged MotB showed fluorescent puncta or spots in the bacterial cell membrane. The pixel intensity of iLOV-tagged MotB was approximately two-fold lower than the intensity of GFP-tagged MotB (**Figure 4.A & B**). In the periplasmic variant (v2), we did not observe fluorescent spots in the cell membrane for either iLOV (**Figure 4.C**) or GFP-tagged MotB (**Supplementary Fig. 9**) and the fluorescence intensity was equivalent to the autofluorescence of the non-fluorescent background strain (**Figure 4.D**). Insertion of short flexible GS linkers removed puncta and decreased the maximum fluorescence intensity inside the regions of interest shown (**Figure 4.E & F**). As a positive control we used monomeric mCherry which has shown brighter fluorescence in the bacterial periplasm than GFP (Pena et al., 2020). Like GFP and iLOV-tagged periplasmic variants (v2), mCherry-tagged MotB were motile but showed limited fluorescence (**Supplementary Fig. 11**).

**Figure 4.**
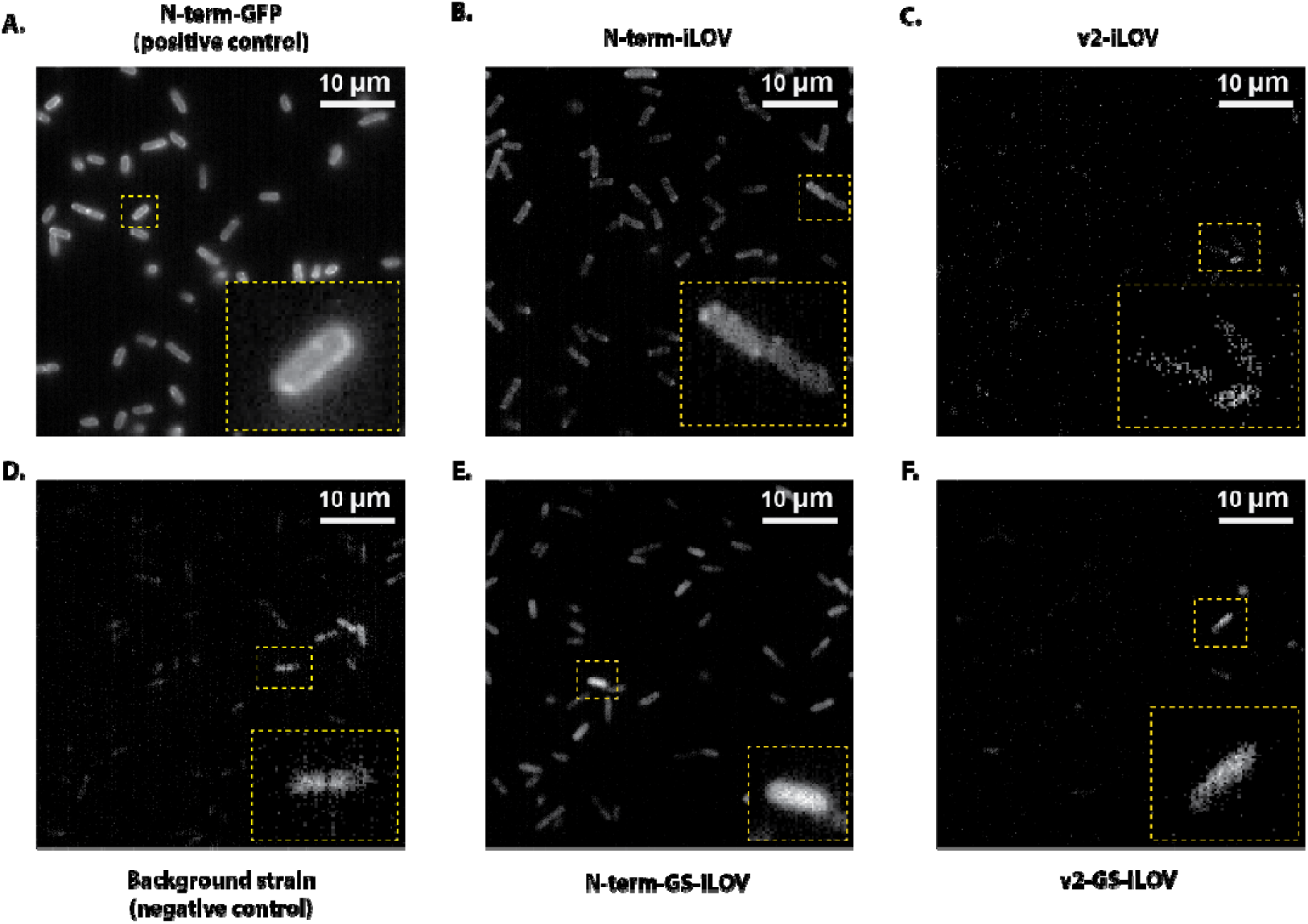
Epifluorescence images of iLOV- and GFP-tagged variants with and without GS linkers in the bacterial membrane. Minimum pixel intensity was set at 110 a.u. for all images. (A) N-terminal GFP-MotB (positive control, maximum – 318 a.u.), and (D) untagged MotB (negative control, maximum – 115 a.u.) (B) N-terminal iLOV-MotB (maximum – 148 a.u.), (C) v2 iLOV-MotB (maximum – 145 a.u.), (E) N-terminal iLOV-MotB-GS (maximum – 104 a.u.), and (F) v2 iLOV-MotB with GS linker (maximum – 113 a.u.). Images are median filtered over 100 frames.

## Discussion

Fluorescently tagged stators have been useful for quantifying stator dynamics in tethered cells (Leake et al., 2006) as well as in response to various loads applied to the flagellar motor (Heo et al., 2017). These constructs are useful for measuring precisely when and how many stators are engaged, pushing on the rotor during operating cycles of the flagellar motor. However, these constructs have been previously shown to affect rotational speed and, more significantly, switching frequency. In this paper, we labelled MotB with the fluorescent proteins GFP and iLOV in an attempt to engineer new constructs with less impacted stator function than existing constructs.

Previous studies have indicated that the disruption of MotB via insertion or deletion, specifically in/near the plug domain, can decrease cell growth or, in some cases, lead to cell death (Hosking et al., 2006). Our constructs with similar insertions (AsLov2) near the plug domain showed growth rates like wild-type bacteria with native stators, implying no adverse impact on plug function. Periplasmic variants (v2) i.e., fluorescent protein (GFP and iLOV) inserted in/near the plug domain were motile and powered flagellar rotation with similar speed compared with the cytoplasmic variants (N-term) both in tethered cell measurements and bead assays. However, these periplasmic variants (v2) may constrain the conformational movements of the plug and reduce the effective transduction of ion flow into torque. Alternatively, the stator binding and unbinding may be affected through interference with the insertion of fluorescent proteins and conformational rearrangements that need to occur for flagellar motor rotation. Interestingly, iLOV in the periplasmic variant (v2) swam faster than the GFP counterpart. GFP with high molecular size of ∼28 kDa compared to iLOV (∼13 kDa) could have impacted the flagellar motor to slow down (Heo et al., 2017). This work is first to test the effect of fluorescent proteins with different molecular sizes in the bacterial motility, thus, we hypothesize that small molecular size of iLOV may have resulted in reduced steric and structural interference affecting stator rotation (Deme et al., 2020; Santiveri et al., 2020). For both cytoplasmic (N-term) and periplasmic variants (v2), inserting short and flexible GS linkers in the iLOV-tagged MotB improved motility as measured by the spread on semi-solid agar plates but not much in tethered cell and bead measurements. Along with bacterial swimming speed, spreading of cell populations in the swim plates depends on growth rate and chemotactic ability (Ha et al., 2014), whereas single cell measurements via bead or tethered cell directly measure torque and speed output of the individual motors. We observed no difference in the growth rate due to presence of linkers for the strains with LOV domain, like iLOV in the stator (**Supplementary Fig. 2**). Similarly in single cell or individual motor, we measured switching properties via bead measurements and observed no effect from the addition of linkers **(Supplementary Fig. 7 & 8)**.

When MotB is tagged with a fluorescent protein it would be expected to fluoresce in the bacterial membrane as a bright puncta, or spots (Heo et al., 2017; Leake et al., 2006; Tipping et al., 2013). Both GFP and iLOV-tagged MotB in the cytoplasmic variant (N-term) displayed puncta, however, the fluorescence intensity of iLOV was lower than that of GFP. This is likely due to weak fluorescence of iLOV compared to GFP (Mukherjee et al., 2013) that was verified by low intensity of iLOV (without MotB fusion) when expressed in bacterial cytoplasm (**Supplementary Fig. 10**). For the periplasmic variant (v2) no distinct puncta were observable either for iLOV or GFP tagged MotB. Limited fluorescence in the bacterial periplasm could arise from impeded folding in the periplasm, or maturation defects (Meiresonne et al., 2019) or the oxidising environment and toxicity due to protein overexpression (Arts et al., 2015; Meiresonne et al., 2017; Wilks & Slonczewski, 2007). Monomeric mCherry, which has been observed to fluoresce with greater intensity in the bacterial periplasm than GFP (Pena et al., 2020) also showed limited fluorescence in the periplasmic variants (v2) (**Supplementary Fig. 11**). Similarly, insertion of GS linkers showed no improvement in the fluorescence of GFP and iLOV for both cytoplasmic (N-term) and periplasmic variants (v2). We speculate that protein folding is disrupted in periplasmic variants (v2). Future works such as using iLOVs with improved brightness (Liang et al., 2022), rigid linkers of varying lengths (Heo et al., 2017) and targeting new sites between plug and PG binding domains of MotB stator protein could generate improved labelled stators for experimental use.

## Conclusion

This paper tested the effect of tagging MotB near the plug domain with fluorescent proteins. Our findings showed that insertion of GFP and iLOV near the plug domain did not disrupt the function of stator complex, but also did not fluoresce brightly and these constructs were not suitable for single molecule measurements of stator binding and unbinding. We have successfully shown that periplasmic sites near the plug are suitable for the addition of either fluorescent reporters or light-switchable domains, but further work is needed to improve both the fluorescence and light-response of these domains to enable direct light-activation of the flagellar stator complex.

## Materials and methods

### Plasmid construction

Four different chimeric constructs of *Escherichia coli* MotB and GFP or LOV protein domains were designed *in silico* and synthesized as gBlocks (IDT) for experimental testing *in vivo* in *E. coli*. All constructs employed in the study were cloned into the pBAD33 vector (chloramphenicol resistant) and expressed using an Arabinose-inducible pBAD expression system (Guzman et al., 1995). To construct plasmids for four variants (v1, v2, v3, and v4), fragments containing AsLOV2 and MotB with two restriction sites were PCR amplified from DNA fragments, then double-digested (SalI and PstI from NEB), ligated (T4 DNA-Ligase, NEB), and transformed in the background strains. Two background strains were used: RP3087 (ΔmotB, for swim plate assay) (Islam et al., 2023) and SYC35 (ΔmotA ΔmotB fliC-sticky) for transformation (Kuwajima, 1988; Sowa et al., 2005). Similarly, restriction-digestion was used to construct plasmids containing iLOV, GFP, and mCherry in the N-terminal and v2 positions of MotB. Quikchange site-directed mutagenesis was employed to insert GS (Glycine-Serine) linkers in these plasmids at the N- and C-terminal flanks of the chimeric insert. The sequences of primers used for restriction-ligation and site-directed mutagenesis are listed in **Supplementary Table 2**.

### Bacterial sample preparation

Frozen aliquots of cells (grown to saturation and stored in 25% glycerol at −75°C) were aerobically grown overnight in 5 mL of Lysogeny Broth (LB, 10 gL−1 bacto tryptone, 5 gL−1 yeast extract, 10 gL−1 NaCl) at 37°C in a continuous shaking incubator at 200rpm. The next day, a new culture was started by diluting 50µL of the overnight culture in 5mL of Tryptone Broth (TB, 10gL−1 bacto-tryptone, 5 gL−1 NaCl) containing 100µM L-arabinose at 30°C. Cells were grown for about 6 hours, shaking at 200rpm, to a final optical density at 600 nm of 0.5 − 0.6. All the culture media contained 100 µgmL−1 ampicillin and 25 µgmL−1 chloramphenicol.

### Swim plate assay

For swim plates, a single colony of bacteria was inoculated in a semi-solid agar medium and incubated at 30°C. To test the effect of blue light in bacteria containing light-responsive domain such as LOV, we built a blue light projector system to illuminate swim plates from the top (Zhang et al., 2020). To build this system, a blue LED of 465 nm (Meccanixity) was installed in the car door light projector (**Supplementary Fig. 3**). The LED was powered by a 0-5 VDC power source. The swim plate inoculated with bacteria was placed 12 cm below the blue LED, and the entire system was placed inside the incubator at 30°C. The swim ring was imaged using a gel doc imager (Bio-Rad) and measured using ImageJ software.

### Tethered cell assay

The rotational speed of cells tethered to the microscope glass slide via a single flagellum was measured to analyse individual flagellar motors. Bacterial cells with short sticky filaments (SYC35 i.e., Δ*motA* Δ*motB fliC*-sticky) where MotA and MotB stator proteins were coexpressed on two different plasmids (**Supplementary Table 2**) were tethered to a glass surface. An inverted phase contrast microscope (Nikon) with a 20X objective lens was used to image the rotation of the tethered cells. 20-second-long videos were acquired with a camera acquisition rate of 20 fps and the rotational speed was measured using LabView software. The number of cells used for the measurements and analysis are listed in the **Supplementary Table 3**.

### Bead assay

The same bacterial strains were used for both the bead assay and the tethered cell assay. To obtain short flagellar stubs, bacteria were mechanically sheared by passing 1 mL of cell culture back and forth between two syringes connected by 21-gauge needles and a thin tube (Gabel & Berg, 2003). Cells were then centrifuged (3000 rpm for 3 min) and resuspended in motility buffer (MB: 10 mM potassium phosphate, 0.1 mM EDTA, 10 mM lactic acid, pH 7.0). Experiments were performed in a ‘tunnel slide’ made of two cover slips (Menzel-Glaser #1.5) separated by a layer of parafilm, with a tunnel cut into it, melted to the coverslips. Two laser-cut holes in the top coverslip allowed fluid exchange. The slides were cleaned with ethanol and sterile water before use. First, 100 µL of poly-L-lysine (Sigma-Aldrich P4707) was introduced into the tunnel slide and incubated for 2 minutes, followed by flushing with 200 µL of MB. Next, 100 µL of cell suspension was flushed through the slide, allowing cells to settle on the coverslip for 10 minutes. Unattached cells were washed out with MB. Finally, a 1/300 dilution of polystyrene beads (1.1 µm in diameter, Sigma-Aldrich) in MB was flushed through. Beads were allowed to spontaneously bind to “sticky” truncated filaments and after about 10 min unattached beads were washed out with MB. Motor rotation was measured by monitoring the rotation of the bead using a custom made bright-field microscope. The sample was illuminated with a 660 nm LED (Thor labs, M660L3) and imaged with a 100 × 1.45-NA objective (Nikon) onto a high-speed CMOS camera (Optronics CL600×2/M). Acquisitions of rotating beads were taken at a sampling rate of 1 kHz.

### Fluorescence assay

An Elyra-7 (Zeiss) microscope was used to test the fluorescence of bacterial strains expressing MotB tagged with GFP and fluorescent LOV domain (iLOV). An oil immersion objective lens with 63 × NA-1.46 was used for all the bacterial strains. Laser light (5 mW) of 488 nm wavelength was used to excite GFP and iLOV. Bacterial samples were prepared as in the tethered cell assay and bead assay to test fluorescence along with rotation. 100 image frames were acquired with an exposure time of 100 ms and an image of size 512 by 512 pixels. A single image was generated from a median filter of the 100 images.

### Data analysis

For the bead assay, the data analysis was performed using custom LabView and Python scripts. The x(t), y(t) positions of the rotating bead were determined by using a cross-correlation analysis of the bead image with a numerically generated kernel pattern (Lipfert et al., 2011). The drift of the circular trajectory was corrected by subtracting a linear interpolation of x and y from their respective raw values. The elliptical trajectories of the beads, assumed to be the projection of a tilted circle, were transformed into circles by stretching the minor axis of the ellipse. Speed was calculated by taking the derivative of the angle, and the speed traces were filtered using a 70 ms window median filter. Switching events were detected by identifying crossings of two thresholds: the positive speed threshold was set at 2/3 of the mean positive speed (Bai et al., 2010; Heo et al., 2017), and the negative threshold was set at 2/3 the mean negative speed, except in cases where the mean negative speed was greater than -3 Hz, in which case the negative threshold was set to -3 Hz. The switching frequency of the motor was calculated as the number of detected switching events divided by the duration of the measurement. MATLAB and Prism GraphPad were used to analyse and plot the data. Statistical analysis was performed using an unpaired *t*-test (except for the swim plate assay, comparing motility in dark and light conditions of the same bacterial strain, where a paired t-test was performed). Data were shown as mean ± standard deviation with error bars. Significance levels while comparing two or more two data sets were presented as p-values.

## Acknowledgements

We thank Peter L Voyvodic for wetlab support at CNRS and Monerh Al-Shahrani for experimental training in DDM at RMIT. M.A.B.B was supported by a Scientia Fellowship from UNSW, by HFSP RGY0072/2 and US Navy Office of Naval Research, Research grant N62909-22-1-2051. A.B.B. and A.L.N. were supported by the French National Research Agency (ANR) PHYBION project grant ANR-23-ERCB-0005-01. A.L.N was further supported by the ANR PHYBABIFO project grant ANR-22-CE30-0034. F.P. was supported by the ANR project grant ANR-23-CE30-0010. The CBS is a member of the France-BioImaging (FBI) and the French Infrastructure for Integrated Structural Biology (FRISBI), two national infrastructures supported by the ANR (ANR-10-INBS-04-01 and ANR-10-INBS-05, respectively).

