## Supplementary Methods 1-2, Supplementary Figures 1-10, Supplementary Tables 1-2. for "Insertion of fluorescent proteins near the plug domain of MotB generates functional stator complex"

**Supplementary Method 1. Differential dynamic microscopy (DDM)**

A single colony was picked up from LB agar streak plates and inoculated in LB broth for overnight culture at 37°C, at 180 rpm in shaking incubator. 350 µL of overnight culture was added to 35 mL of TB broth and incubated at 30°C in a shaking incubator for 4-5 hours till the OD600 reaches to 0.5-0.6. Then, it was diluted by 5 times by adding 5 mL of PBS to 1 mL of subculture. Then, 50-70 µL of the diluted sample (OD600 around 0.1 -0.2) was loaded in a rectangular capillary tube (VitroTubes™), sealed with vaseline on both ends of the capillary and imaged with an Olympus IX-71 inverted microscope with 10x phase contrast and Köhler illumination. Samples were imaged in the middle plane of the capillary tube. Videos of 1 minute at 100 fps (i.e., 6000 frames) were acquired at a frame size of 512 x 512 pixels using a Mikrotron MC-1362 camera connected to a frame grabber card (Euresys Grablink). LabView code was used to process the video, followed by using in house MATLAB codes for the analysis and data visualisation (Wilson et al., 2011). The mean swim velocity for each bacterial strain was obtained by fitting the DDM parameters versus q-values which is explained in detail in the review paper (Al-Shahrani & Bryant, 2022).

**Supplementary Method 2. Corrected total cell fluorescence (CTCF)**

To quantify the fluorescence signal in bacterial cytoplasm, the corrected total cell fluorescence (CTCF) value was calculated. First, a region of interest (ROI) was drawn around whole bacterial cell, then, values for 1) $I_D$, the integrated density (the sum of pixels in the region of interest), 2) $A_{ROI}$, the area of the ROI, and 3) $F_B$, the mean fluorescence of background pixels were measured (as determined by dark pixels outside the boundary of the cell). Finally, CTCF was calculated in ImageJ by using the formula:

$$\text{CTCF} = I_D - (A_{ROI} \times F_B).$$

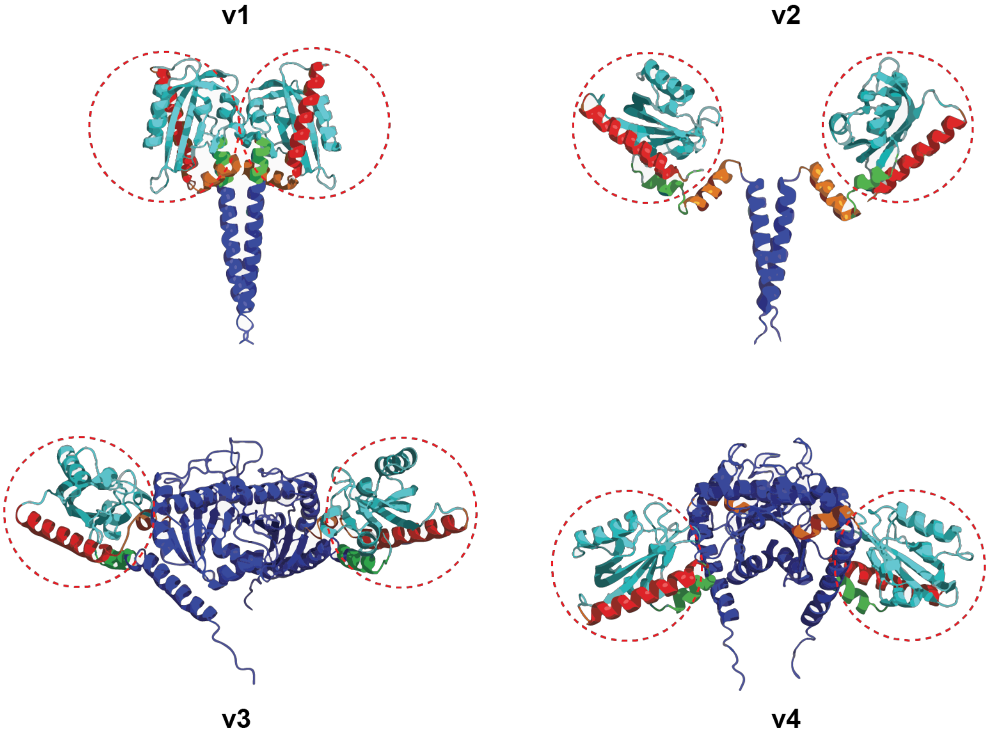


**Supplementary Fig. 1. AlphaFold structure of AsLOV2 domain inserted in MotB.** v1 and v2 represented the insertion of the LOV domain before and after the plug domain (dark orange) respectively. v3 and v4 represented the insertion of the LOV domain before and after the short α2-helix of the PG Binding domain (dark orange) respectively. AsLOV2 (LOV domain) is marked within a dotted circle with A'-α-helix (green), LOV core (light blue), and C-terminal J-α-helix (red). Cryo-EM structure of N-terminal of MotB (blue-colored) with plug domain (orange-colored) of Campylobacter jejuni (PDBID: 6ykm). X-ray crystallography structure of MotB PG binding domain (blue-colored) from Salmonella enterica (PDBID: 2zvz). X-ray crystallography structure of AsLOV2 from Avena sativa in a dark state (PDBID: 2v1a). AlphaFold was used for the combined structure of the AsLOV2 domain integrated with MotB (plug domain and PG binding domain).

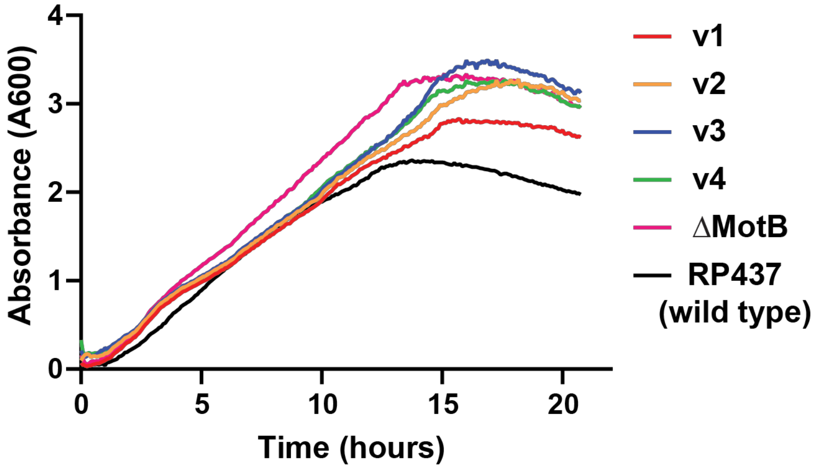


**Supplementary Fig. 2. Line graph of growth curves, recorded as absorbance at 600 nm, incubated at 37°C for 24 hours.** Bacterial strains: MotB tagged with AsLOV2 (v1, v2, v3, v4), MotB deleted strain as background strain (ΔMotB), and wild-type motile strain as positive control (RP437). Absorbance was recorded every 15 minutes at time intervals for 24 hours. Average absorbance was plotted versus time, taken from four replicates for each bacterial strain

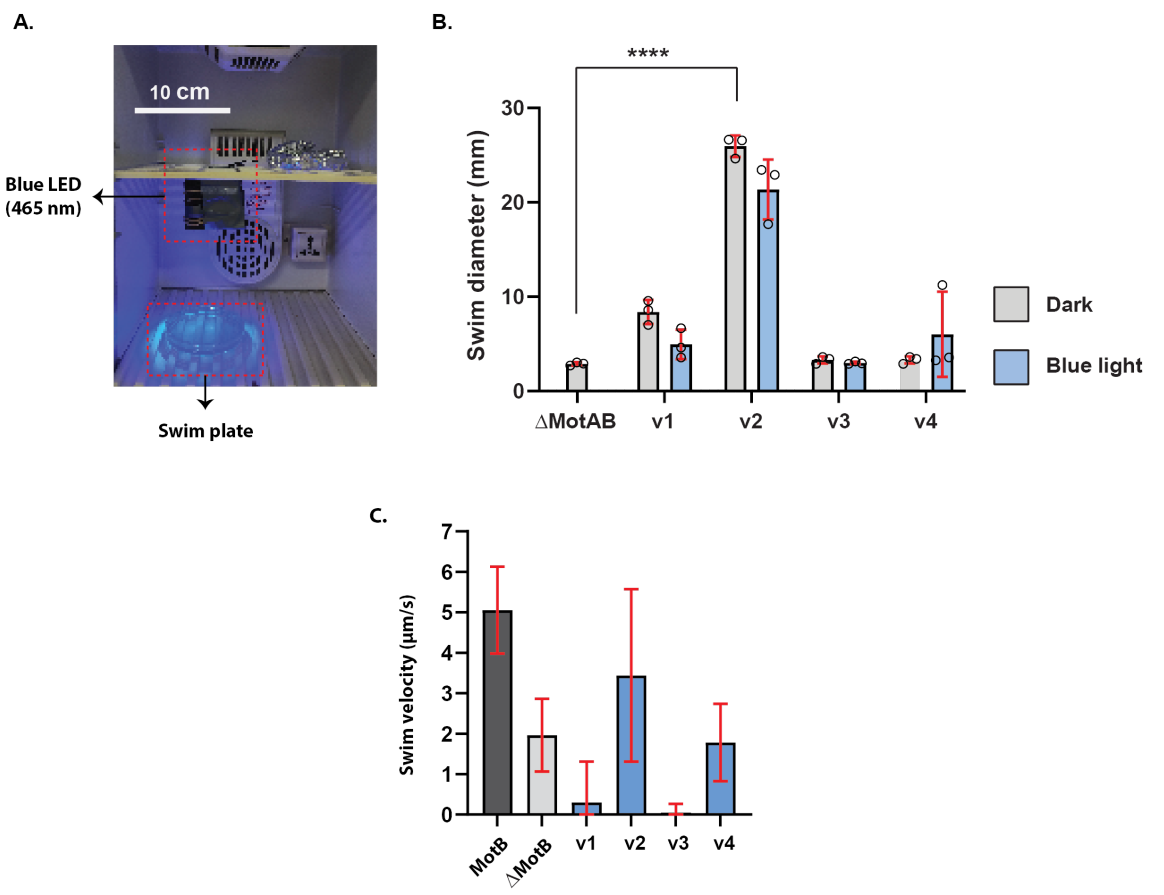


**Supplementary Fig. 3. Effect of blue light (465 nm, 150 lux intensity) in the motility of bacteria with LOV inserted MotB.** Bacterial strains; ΔMotAB - non-motile, LOV domain inserted MotB; v1 (before plug domain), v2 (after plug domain), v3 (before PG domain), and v4 (after PG domain). Blue-coloured bar – blue light illumination and light-grey coloured – dark condition. (A) Blue light illumination system (scale bar of 10 cm). The swim plate was placed under the blue LED projector (465 nm) at a distance of ~ 12 cm inside an incubator. The swim plate and blue LED are labelled and denoted by a dotted red rectangle in the figure above. The intensity of blue light was measured at ~150 lux by lux meter. (B) Bar graph of swim ring diameter (mean ± standard deviation, triplicates data) measured from swim plate assay. (C) Swimming velocity of bacterial strains in liquid media measured by differential dynamic microscopy (DDM).

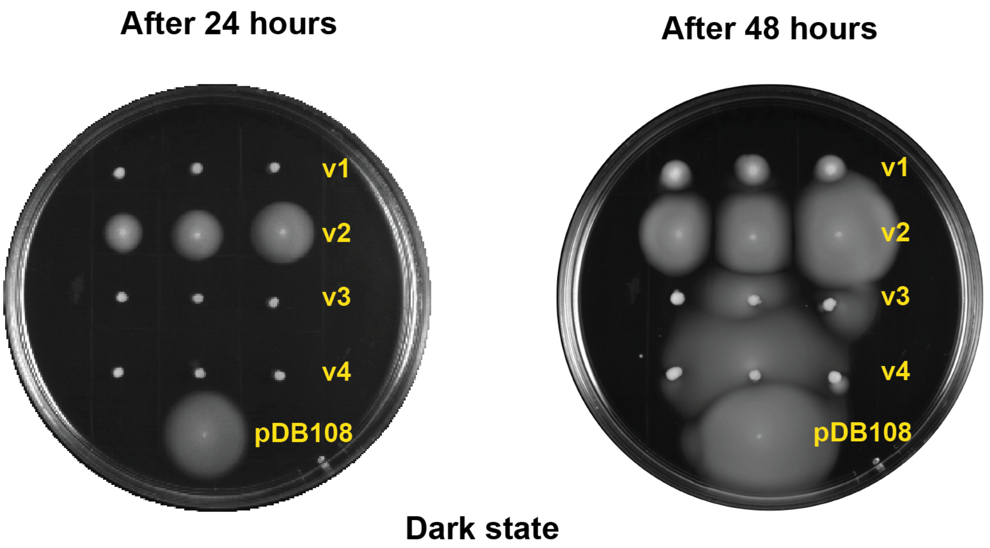


**Supplementary Fig. 4. Image of swim plates after 24 hours (left) and 48 hours (right) of incubation at 30°C. Bacterial strains:** MotB tagged with AsLOV2 (v1, v2, v3, v4), and wild-type motile strain as a positive control (pDB108 – $MotA_5MotB_2$ expressing bacterial strain). Three colonies for each bacterial strain were inoculated in a swim plate (except for pDB108).

.

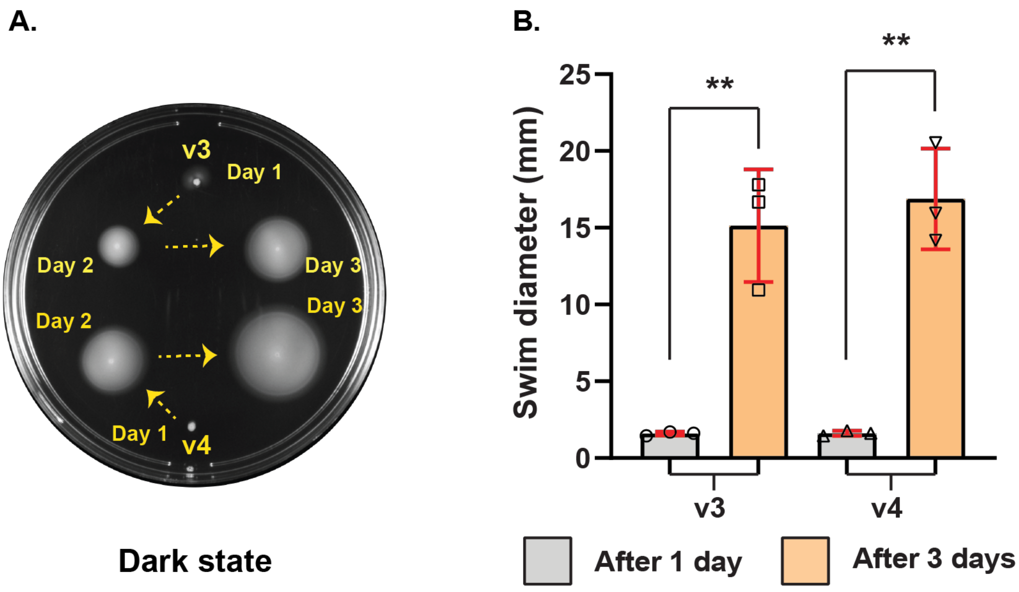


**Supplementary Fig. 5. Swim plate assay depicting the motility status of v3 and v4.** (A). Image of swim plate showing the motility of colonies after incubating for 1 day, 2 days, and 3 days at 30°C. (B). Bar graph of swim diameter (mean ± standard deviation) of v3 and v4 measured for colonies incubated for 1 day and 3 days at 30°C.

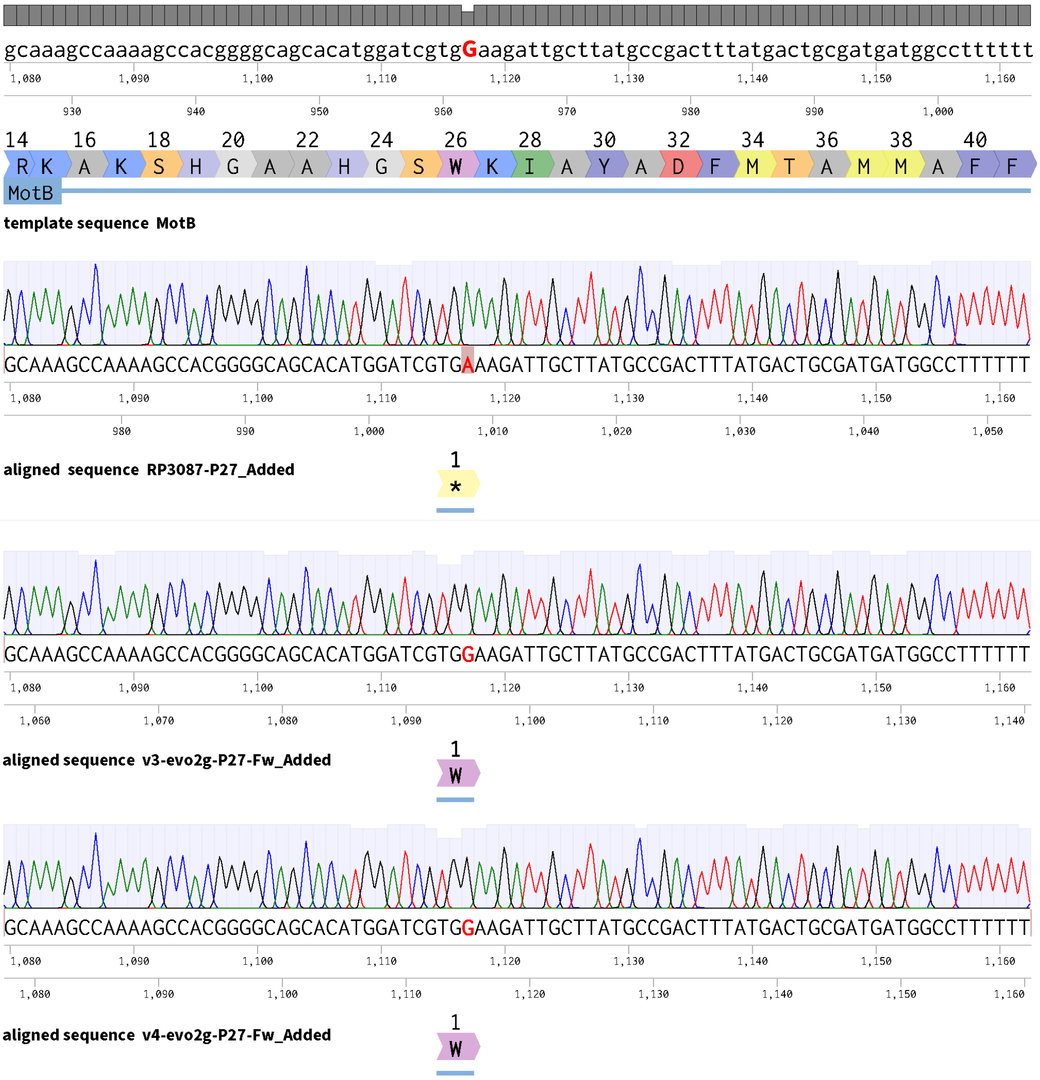


**Supplementary Fig. 6. Sanger sequencing result of motB gene fragment amplified from the bacterial genome.** In RP3087 i.e., ΔMotB background bacterial strain, $26^{th}$ amino acid tryptophan (**W**) was converted to stop codon (*****) by single nucleotide mutation (TG**G** to TG**A**) to stop the translation of MotB. After 2-3 days of incubation in a swim plate at 30°C, variants v3 and v4 (contains plasmid expressing LOV domain inserted at PG binding domain of MotB in ΔMotB background strain) rescued its motility by reverting this mutation from TG**A** to TG**G** (i.e., from stop codon '*' to tryptophan '**W**').

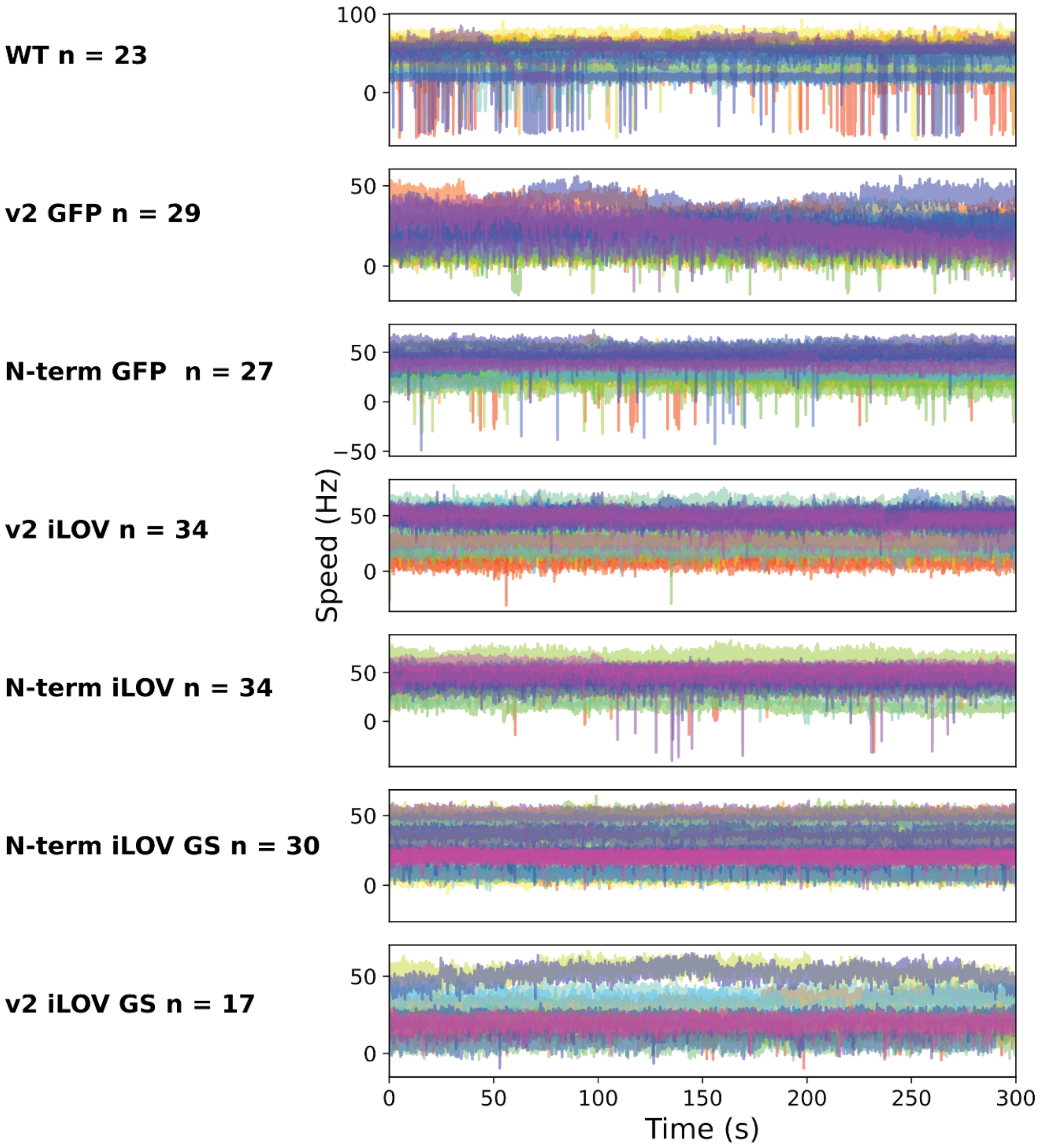


**Supplementary Fig. 7. Motor speed traces of individual CCW-biased motors rotating 1.1 µm beads (Left) and probability distributions of motor speeds (Right) (without GS linkers).** Positive speeds indicate counterclockwise (CCW) rotation, while negative speeds indicate clockwise (CW) rotation. The blue colour represents CCW-biased motors, and the red colour represents CW-biased motors. Histograms correspond to individual motor measurements, and thick lines denote the average across all measurements. The number of motors measured is 23 for WT, 29 for v2 GFP, 27 for N-term GFP, 34 for v2-iLOV and 34 for N-term iLOV.

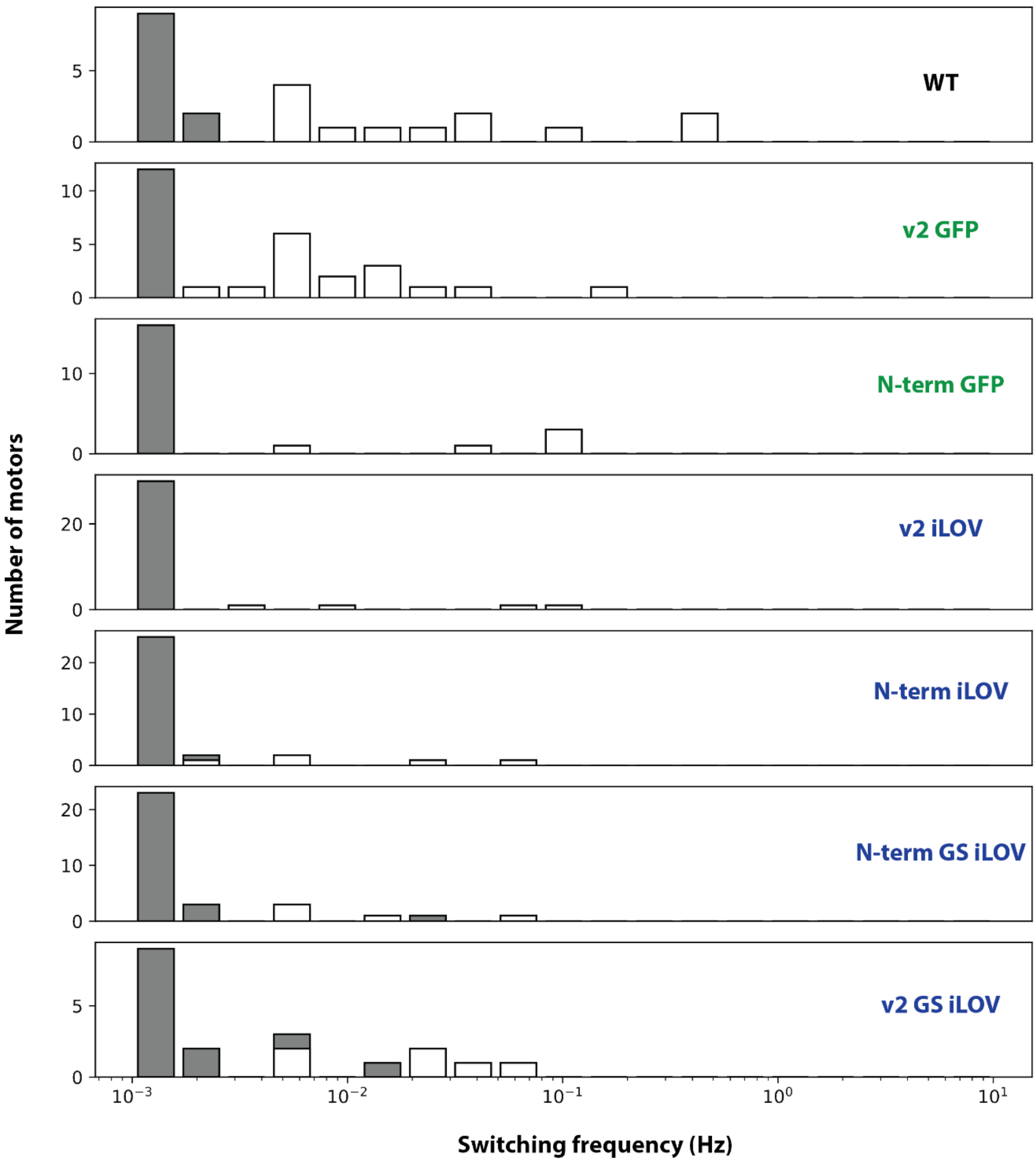


**Supplementary Fig. 8.** Switching frequency distributions plotted for iLOV and GFP tagged MotB in N-terminal and v2 position from all the individual speed traces of (n) number of motors (Supplementary Fig. 7). Grey bar denotes the motors that did not switch during the measurements. The number of motors (n) measured was 23 for WT, 29 for v2 GFP, 27 for N-terminal GFP, 34 for v2-iLOV, 34 for N-terminal iLOV, 30 for N-terminal iLOV GS, and 17 for v2 iLOV GS. Here, 'GS' indicates the linker with two amino acids of Glycine-Serine.

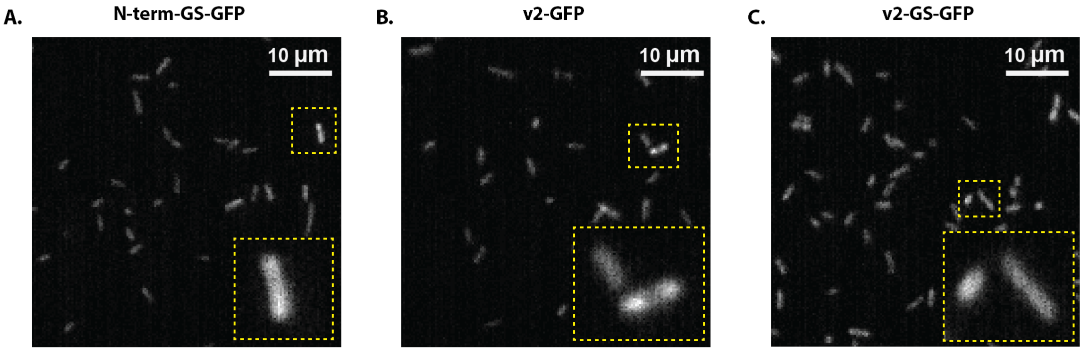


**Supplementary Fig. 9. Fluorescence images (median of 100 frames) of GFP tagged in MotB (cytoplasmic and periplasmic end with and without GS linkers) in the bacterial membrane.** Minimum pixel intensity was set at 110 a.u. (arbitrary unit) for all images. (A) MotA + GFP-MotB-GS (maximum – 142 a.u.), (B) MotA + MotB-GFP-v2 (maximum – 145 a.u.), and (C) MotA + MotB-GFP-v2-GS (maximum – 138 a.u.). 100 frames of fluorescence images were recorded in EPI mode of illumination, excited by a 488 nm laser of 5 mW power and 100 ms of exposure time. All intensity measurements were captured by an sCMOS camera.

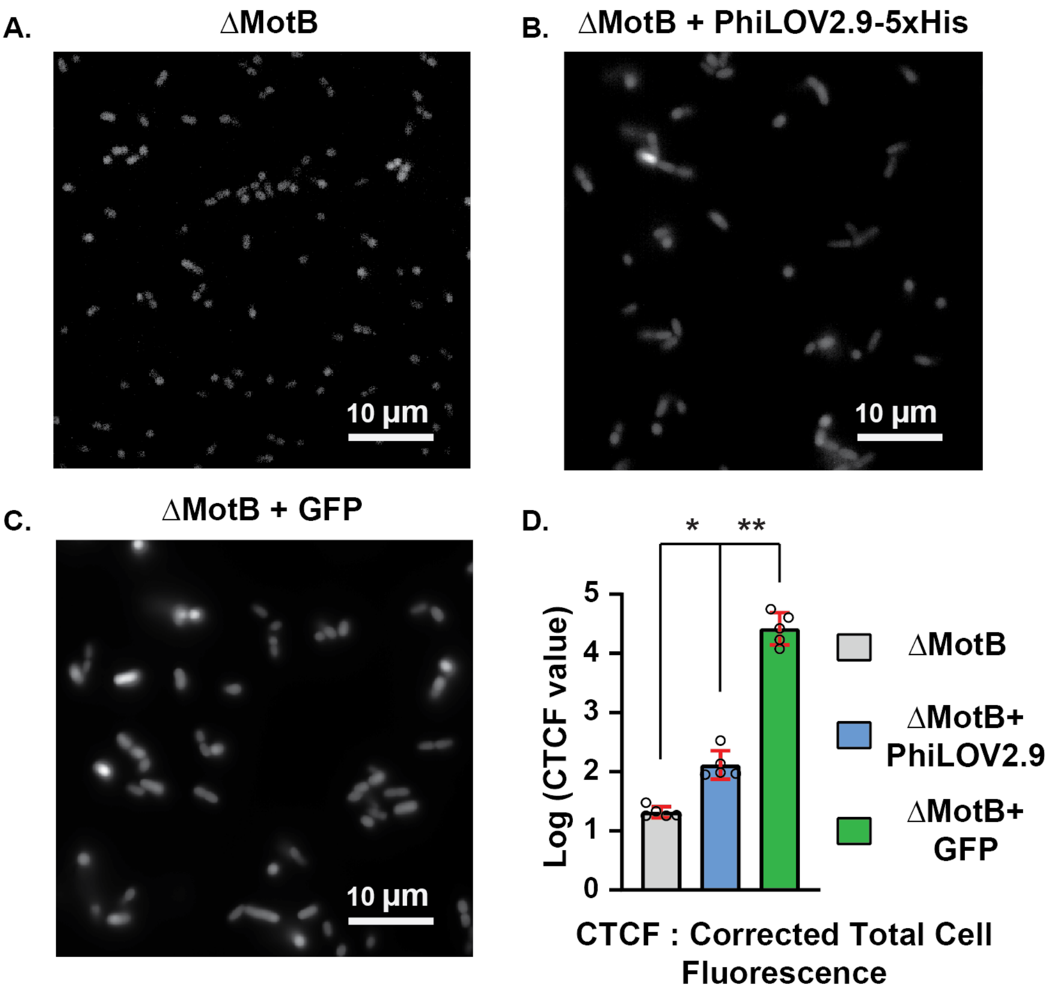


**Supplementary Fig. 10. Fluorescence of cytoplasmic expression of ILOV and GFP fluorescent proteins.** The minimum pixel intensity was set at 100 for all three images. Fluorescence images (median of 100 frames) of (A) ΔMotB (maximum - 145), (B) ΔMotB + ILOV-5xHis (maximum – 342), and (C) ΔMotB + GFP (maximum - 29357). (D) Fluorescence measurements using corrected total cell fluorescence (CTCF) value. Five cells for each bacterial strain were used to calculate the average CTCF value (**Error! Reference source not found.**) and plotted as a logarithmic value. All intensity measurements were captured by an sCMOS camera.

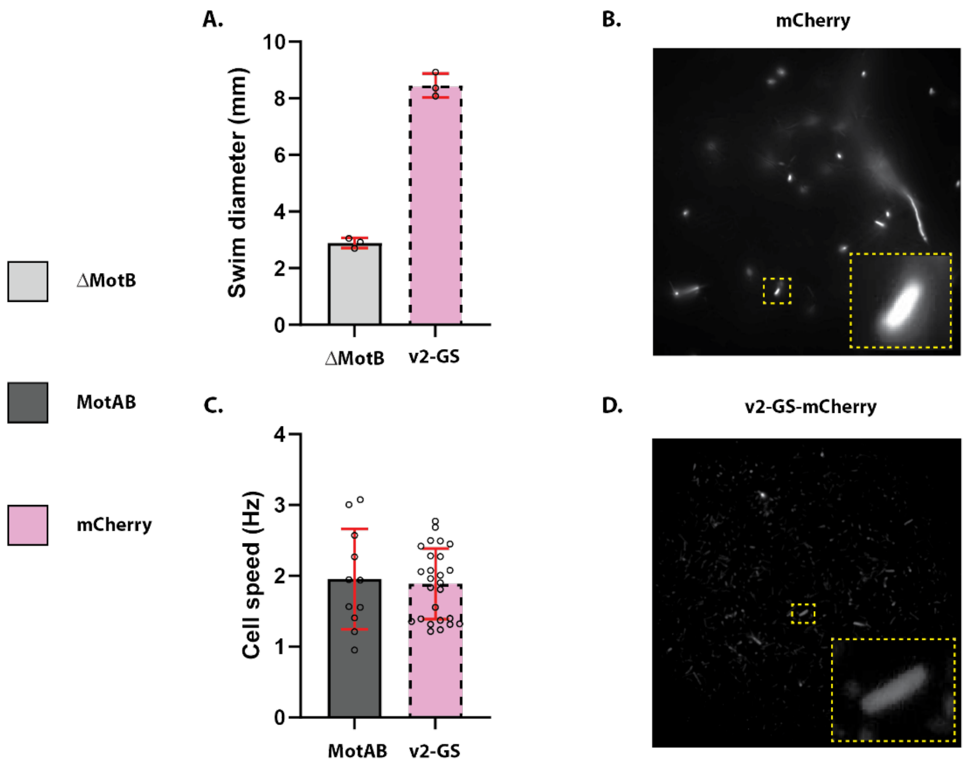


**Supplementary Fig. 11. Motility and fluorescence of mCherry-v2-GS bacterial strain.** (A) Bar graph of swim ring diameter (mean ± standard deviation, triplicates data) measured for the bacterial strains tested (B) Bar graph of rotational speed (mean ± standard deviation) for the bacterial strains tested. (C) Cytoplasmic expression of mCherry (positive control) (D) MotA + MotB-GFP-v2-GS. 100 frames of fluorescence images were recorded in EPI mode of illumination, excited by a 488 nm laser of 5 mW power and 100 ms of exposure time. All intensity measurements were captured by an sCMOS camera.

**Supplementary Table 1.** List of the average swim velocity (mean ± with standard deviation) from differential dynamic microscopy (DDM).

| ***E. coli* strain** | **Condition** | **Plasmids** | **Speed velocity (µm/s)** |
|---|---|---|---|
| RP3087 | Dark | pMotB | 5.1 ± 1.1 |
| RP3087 | Dark | -- | 2.0 ± 0.1 |
| RP3087 | Dark | pMotB-LOV-v1 | 0.3 ± 1.0 |
| RP3087 | Dark | pMotB-LOV-v2 | 3.4 ± 2.1 |
| RP3087 | Dark | pMotB-LOV-v3 | 0.1 ± 0.2 |
| RP3087 | Dark | pMotB-LOV-v4 | 1.8 ± 1.0 |

**Supplementary Table 2.** List of the average rotational speed of tethered cell (mean ± with standard deviation) from tethered cell assay.

| **Primers** | **Sequence** |
|---|---|
| **Restriction-Digestion** | |
| P20 | CAGTGAATGGGGGTAAAT |
| P24 | GGTTGGACTCAAGACGATAG |
| **Site-directed mutagenesis** | |
| GFP-MotB link Fw | GGCAGCATGAAAAATCAGGCTCACC |
| GFP-MotB link Rv | CTTGTACAGTTCGTCCATG |
| iLOV-MotB link Fw | GGCAGCATGAAGAATCAAGCGCATC |
| iLOV-MotB link Rv | TTTATCATCATCATCTTTATAATCGCT |
| v2-EGFP-909a | GGATGAGCTTTACAAGGGCAGCCCCCTTGCTACCGC |
| v2-EGFP-192b | TCCCCTTTACTCACCATGCTGCCCGTACGGAAGTATTCGG |

GGCAGC: GS linker nucleotide sequence

**Supplementary Table 3.** List of the average rotational speed of tethered cell (mean ± with standard deviation) from tethered cell assay.

| ***E. coli* strain** | **Plasmids** | **No. of cells (n)** | **Rotational speed (Hz)** |
|---|---|---|---|
| SYC35 | pMotA, pMotB | 11 | 2.0 ± 0.7 |
| SYC35 | pMotA, pGFP-MotB | 14 | 1.6 ± 0.6 |
| SYC35 | pMotA, pGFP-MotB-GS | 15 | 1.9 ± 0.7 |
| SYC35 | pMotA, pMotB-GFP-v2 | 21 | 2.0 ± 0.6 |
| SYC35 | pMotA, pMotB-GFP-v2-GS | 17 | 1.9 ± 0.7 |
| SYC35 | pMotA, piLOV-MotB | 38 | 1.8 ± 0.7 |
| SYC35 | pMotA, piLOV-MotB-GS | 21 | 2.4 ± 0.6 |
| SYC35 | pMotA, pMotB-iLOV-v2 | 30 | 1.8 ± 0.5 |
| SYC35 | pMotA, pMotB-iLOV-v2-GS | 22 | 1.8 ± 0.6 |
